# An *in silico* analysis of robust but fragile gene regulation links enhancer length to robustness

**DOI:** 10.1101/677641

**Authors:** Kenneth A Barr, John Reinitz, Ovidiu Radulescu

**Affiliations:** Dept. of Human Genetics, University of Chicago, Chicago, IL, USA; Depts. of Statistics, Ecology & Evolution, Molecular Genetics & Cell Biology, University of Chicago, Chicago, IL, USA; LPHI UMR CNRS 5235, University of Montpellier, Montpellier, France

## Abstract

1

Organisms must ensure that expression of genes is directed to the appropriate tissues at the correct times, while simultaneously ensuring that these gene regulatory systems are robust to perturbation. This idea is captured by a mathematical concept called *r*-robustness, which says that a system is robust to a perturbation in up to *r -* 1 randomly chosen parameters. In this work we use this idea to investigate the robustness of gene regulation using a sequence level model of the *Drosophila melanogaster* gene *even-skipped*. We find that gene regulation can be remarkably sensitive to changes in transcription factor concentrations at the boundaries of expression features, while it is robust to perturbation elsewhere. We also find that the length of sequence used to control an expression feature correlates negatively with the number of nucleotides that are sensitive to mutation in both natural and *in silico* predicted enhancers. In all cases, the exact degree of robustness obtained is dependent not only on DNA sequence, but also on the local concentration of regulatory factors. By analyzing both natural and synthetic sequences, we provide strong quantitative evidence that increased sequence length makes gene regulatory systems more robust to genetic perturbation.

**Author Summary:** Robustness assures that organisms can survive when faced with unpredictable environments or genetic mutations. In this work, we characterize the robustness of gene regulation using an experimentally validated model of the regulation of the *Drosophila* gene *even-skipped*. We use a mathematically precise definition of robustness that allows us to make quantitative comparisons of robustness between different genetic sequences or between different nuclei. From this analysis, we found that genetic sequences that were not previously known to be important for gene regulation reduce sensitivity to genetic perturbation. In contrast, we found that gene regulation can be very sensitive to the levels of regulators. This extreme sensitivity was only observed at the boundaries of expression features, where switch-like behavior is desirable. This highlights the importance of considering context when assessing robustness.

## 3 Introduction

Biological systems must be robust to perturbations, both environmental and genetic, in order to maintain their functions in fluctuating circumstances[1]. Some of their robustness properties are general, while others are limited to living systems. Robustness of general cybernetic systems was studied mathematically by von Neumann [2]. von Neumann employed multiplexing and majority rule with Boolean automata, an approach that captured buffering by error control and variation, but failed to treat the flexibility and possibility of adaptation underlying biological robustness. Many authors tried to describe the peculiarities of biological robustness using metaphors such as robust-yet-fragile [3], René’s Thom theory of catastrophes [4, 5] or control theory [6]. The relation between robustness, redundancy and complexity is particularly relevant to biological systems. It has been suggested, that, contrary to the common belief that simple systems are robust, a certain degree of complexity can also lead to stable behavior [7, 8]. This property follows from the very general mathematical principle of measure concentration in high dimension and was first discussed in [7]. The law of large numbers is an instance of this principle, ensuring that non-correlated variation of additive effects is buffered and vanishes when the number of elements increases. Within the same formalism, the concept “robust yet fragile” is made precise by the idea of *r*-robustness: a functional property can be stable with respect to perturbation of up to *r -* 1 randoml chosen parameters and sensitive when *r* parameters are varied simultaneously [7]. Robust biological systems include organismal development, where a form of robustness called canalization assures that all individuals arrive at the same phenotype despite individual genetic variation[9, 10, 11]. Genetic and signal transduction network models have provided mechanistic explanations of developmental robustness and canalization in *Drosophila* [12, 13], *C. elegans* [14], and *S. purpuratus* [15].

In gene networks, the connections between elements represent regulatory interactions which control levels of expression of genes through *cis*-regulatory elements, typically 500 to 1000 basepairs (bp) in length, called enhancers[16]. These sequences contain clusters[17, 18] of binding sites for transcription factors (TFs) that act in combination to direct gene expression in specific spatial domains or tissues. While it is well understood how the dynamics of developmental networks confer robustness to the system, it is poorly understood how or if the enhancers that control these networks contribute to robustness through their organization. An important property of enhancers is their redundancy, which is seen at two levels. The clusters of binding sites in an enhancer typically include multiple binding sites for the same TF, conferring a many-to-one relationship between TF binding sites and gene expression[19]. At a higher level, multiple enhancers can control expression in the same expression domain or tissue type [20], and such “shadow” enhancers are known to increase robustness [21, 22, 23, 24, 25, 26]. These specific experimental findings have remained largely unaddressed at the theoretical level.

Previously described data driven and experimentally well tested models of *Drosophila* development are an ideal system for the theoretical study of robustness. Confocal microscopy has been used to generate spatial and temporal atlases of protein and mRNA levels at single nucleus resolution during the first 4 hours of development[27, 28, 29, 30, 31, 32]. These data provide the basis of sequence level models of gene regulation, which predict gene expression levels as a function of protein levels and DNA sequence [33, 34, 35, 36, 37, 38, 39, 40, 41, 42]. Using such models, it is possible to address how the general principles of gene regulation can confer robustness to enhancers.

In this work we use a previously reported model of gene regulation [41] to model the robustness of the *Drosophila even-skipped* (*eve*) locus with respect to variation in both TF concentration and DNA sequence. This model is described fully in the Appendix to this work and it main features are listed at the beginning of Section 4.2. *eve* codes for the homeodomain protein Eve, whose expression forms seven sharply located stripes, necessary for the formation of parasegments during embryonic development [43]. We specifically assess two types of robustness: distributed robustness and *r*-robustness[7]. We find that gene regulation of *eve* can be extraordinarily sensitive to certain changes in TF concentrations, a property that may help form sharp borders in expression domains. We also find that this gene regulation is *r*-robust with respect to sequence mutation. Expression of *eve* is sensitive only to changes in a few nucleotides of the enhancer. Finally, we show that expression from longer enhancers is sensitive to changes in fewer nucleotides in both natural and *in silico* generated enhancers, indicating that enhancer length confers robustness to genetic perturbation.

## 4 Results

### 4.1 Distinguishing types of robustness

Distributed and *r*-robustness arise in complex systems whose properties depend on a large number of parameters. In the former case, the effect of a single perturbation is small and grows very slowly with the number and size of perturbations. In the latter, weaker case, the system is insensitive to the majority of perturbations, excepting the perturbation of a few sensitive parameters. Gorban and Radulescu[7] formalized these types of robustness and investigated the robustness of a well described signaling pathway. In this work we follow the definitions laid out in Gorban and Radulescu [7, Eqs. 1-2]. We consider the robustness of a positive quantitative property *M* that depends on *n* positive parameters *K* = (*K*_1_, *K*_2_, …, *K*_*n*_), namely *M* = *f* (*K*_1_, *K*_2_, …, *K*_*n*_). The property *M* is *distributedly* robust with respect to changes in these parameters if the variance in *M* is reduced compared to the variance in the parameters *K*, when the parameters are subjected to independent perturbations. That is, considering variance in all independent parameters Var(log *K*_*i*_) = Var(log *K*), *i* ∈ *{*1, …, *n}*, then we consider *M* to be *distributedly* robust if

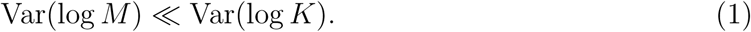

Similarly, if we consider a subset of *r* parameters *I*_*r*_ = *{i*_1_, *i*_2_, …, *i*_*r*_*}* ⊂ 2^*{*1,^ … ^,*n}*^, which we multiply by positive, independent, identically distributed, random scales (*s*_1_, *s*_2_, …, *s*_*r*_), we define *M* as *r*-robust if, given *r* and randomly chosen *I*_*r*_,

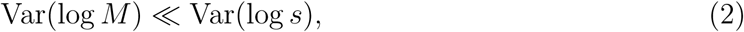

where Var(log *s*) is the variance of each log *s*_*i*_, 1 ≤ *i* ≤ *r*. According to this definition, a property *M* can be *r*-robust only for some values of *r*; typically, it can be robust for small values of *r* and lose this property for large *r*.

To distinguish between these types of robustness, it is useful to study the relationship between the variance of parameters and the variance in the output, or similarly to observe the variance in the output given the number of parameters perturbed. For instance, consider a system and a property *M* that is *r*-robust. In this system, there are *n* parameters, of which *n*_0_ parameters are individually sensitive to perturbation, meaning that *M* is sensibly affected by the perturbation of each of these parameters. If we select *r* of *n* parameters at random, the probability we did not select a sensitive parameter is (1 *- n*_0_*/n*)^*r*^. Then the probability that at least one sensitive parameter was selected is 1 *-* (1 *- n*_0_*/n*)^*r*^. If a sensitive mutation contributes *V*_0_ to the variance, and the effect is not cumulative, then the variance in *M* with respect to *r* mutations is given by

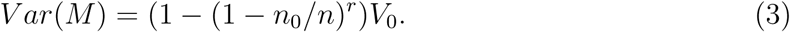

Although *V*_0_ is large, *V ar*(*M*) is small for small *r* and becomes *V*_0_ only when *r* is large enough. The cross-over value of *r* characterizing the loss of robustness is smaller for a large number of sensitive parameters *n*_0_. In particular, the relation (3) can be used to indirectly identify *n*_0_. As discussed in [7], the heterogeneity of the sensitivity of parameters of biochemical systems results from the existence of widely distributed time and concentration scales, a property called “multiscaleness”. In multiscale systems some parameters are important and dominate the others, while a majority of parameters have small effect and play a more static role. The term “sloppy-sensitivity” is sometimes used to designate this situation [44].

In contrast, consider the function *M* = (*K*_1_ + *K*_2_ + …+ *K*_*n*_)*/n*. If all parameters have a variance of *V*_*K*_, then the variance of *M* with respect to *r* will be

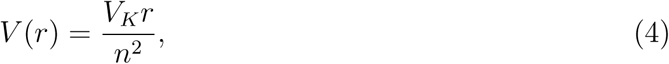

where *r* is the number of parameters that have been independently perturbed. This function grows linearly with respect to *r*. If all parameters are perturbed, the variance of *M* is simply

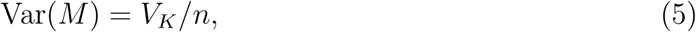

and the maximum variance Var(*M*) is a vanishingly small fraction of *V*_*K*_ when *n* → ∞. This equation illustrates distributed robustness. While the variance of the arithmetic mean scales like 1*/n*, some other robust functions have faster variance decrease with *n*. General distributed robustness is related to concentration of measure in high-dimensional spaces, a phenomenon well known in mathematics. The distribution of a function *f* of *n* variables “concentrates”, meaning that it has vanishing variance when *n* is very large if the function depends on all of the variables but not particularly strongly on one of them (the function should have the Lipshitz property, for details and other examples see Gorban and Radulescu [7]).

In the cases of *r*-robustness or distributed robustness we can assign a number that describes the robustness of the system. When there are sensitive parameters, *n*_0_ describes the robustness of the system, and systems with lower *n*_0_ are more robust. For distributed systems, robustness is described by the ratio of variance of the input to variance of the output, a parameter we call *ρ*. For much of this work we will be working with variables of unknown scale. In light of this, we observe the variance in the fold-change of the input and the fold-change of the output. The fold-change variations are well captured by the variance of the logarithm. More precisely,

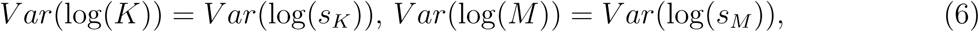

where *s*_*K*_ = *K/K*^′^, *s*_*M*_ = *M/M* ′are the fold changes of *K* and *M* with respect to the reference values *K′*and *M′*respectively. Thus, the robustness ratio *ρ* is given by

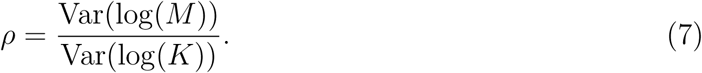

For *r*-robust properties we expect that *ρ* depends on the number *r* of perturbed targets according to

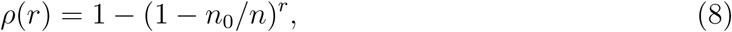

where *n*_0_ is the number of sensitive parameters.

### 4.2 Robustness of mRNA levels with respect to transcription factor concentration

Enhancers interpret the local concentration of transcription factors in order to specify appropriate production of mRNA. In order to determine the degree to which known regulatory mechanisms acting on enhancers contribute robustness to fluctuations in TF concentration, we utilized a model that simulates the regulation of *Drosophila eve* [41], which is expressed in seven transverse stripes across developing *Drosophila* embryos. This model incorporates several mechanisms. The binding of TFs to DNA, including the effects of steric competition and cooperativity, is treated by thermodynamics [45]. Other mechanisms, described phenomenologically, are short-range quenching of transcriptional activators[46, 47, 48], and coactivation of repressors[49, 50, 38]. The functional roles of the TFs used in the model are known from independent experiments, and expression is calculated by summing the bound activators after accounting for the effects of the mechanisms listed above and passing the resulting net activation *N* through a diffusion limited Arrhenius rate law, taking into account competition for interaction with the basal transcriptional machinery. In the study cited above, the model is able to accurately treat the expression pattern of stripes between 35.5% and 92.5% embryo length (1A), identify the enhancers within the *eve* regulatory locus, and simulate the effects of mutations to the environment in *trans*. A detailed description of the model is provided in the Appendix.

To test how this system responds to fluctuations in TF levels, we simulated changes in TF levels by multiplying them by the fold ratio

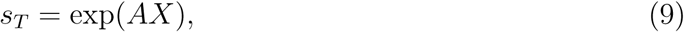

where *A* is a parameter that sets the size of fluctuations, and *X* is a random number drawn from a uniform distribution between −1 and 1. Because TF levels are in arbitrary units, we observe the fold change in TF levels and the fold change in resulting mRNA and computed the ratio *ρ* (Eq 7) after simulating 10,000 fluctuations.

We find that sensitivity to fluctuations in individual TFs varies with respect to position in the embryo. For instance, if we observe sensitivity to fluctuations in the TF Giant (Gt) at the interstripes, borders, and peak of the second *eve* stripe (positions indicated in Fig 1A), we find that at the anterior interstripe and border, expression is not robust to changes in Gt regardless of the magnitude of fluctuation (Fig 1B). This embodies the well established fact that Gt controls the anterior border of *eve* stripe 2 [51, 19, 52]. In contrast, the posterior interstripe of stripe 2 is insensitive to fluctuations in Gt. Notably, at the peak of stripe 2, expression is robust against small fluctuations and sensitive to large ones (Fig 1B).

**Figure 1.**
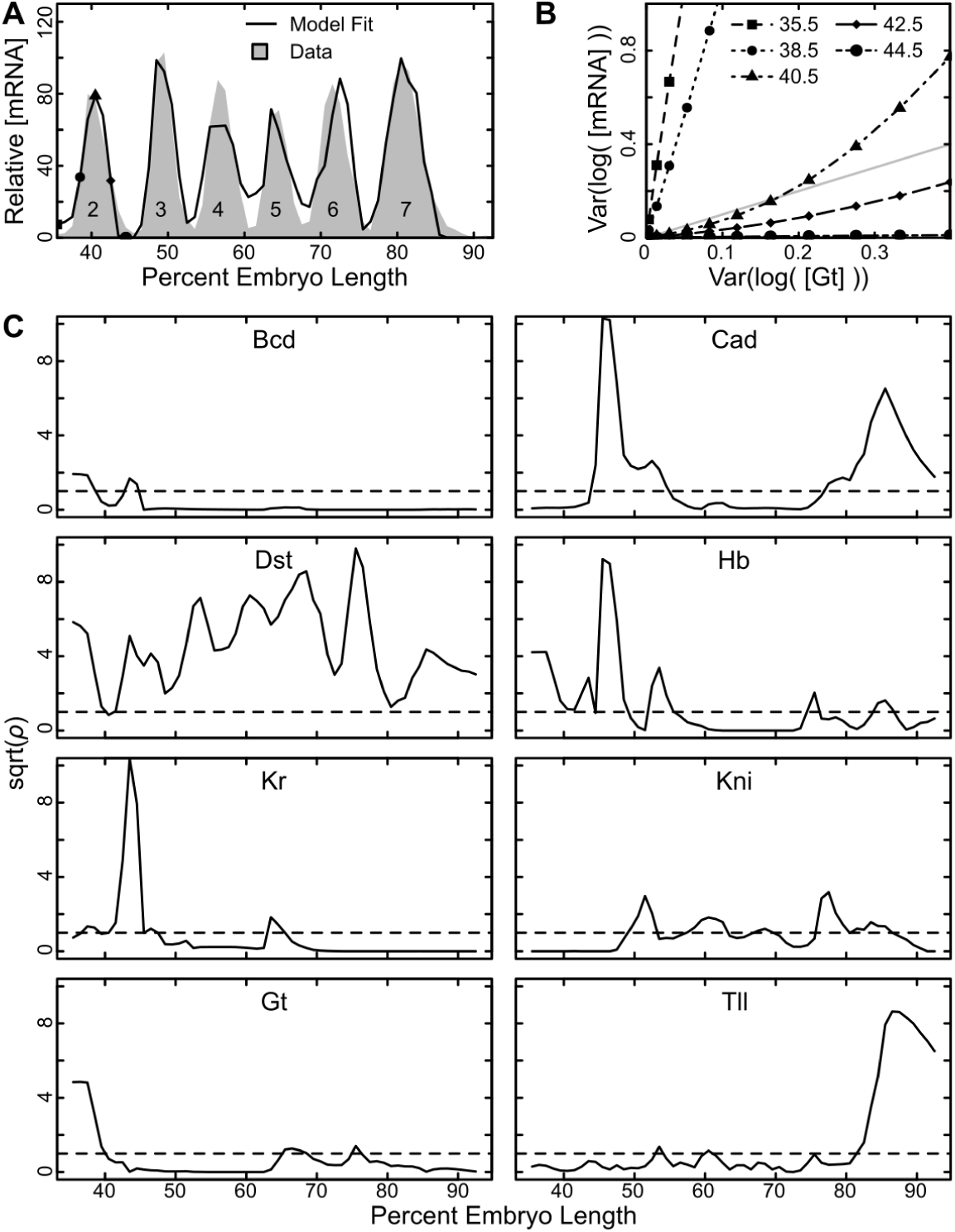
Robustness of *eve* expression to variation in TF concentration. (A) The relative expression of *eve* mRNA along a 10% wide strip along the anterior posterior axis (gray shading) and the model fit to the same data (black line). *eve* stripe number is indicated. (B) The relationship between variation in fold-change TF concentration and fold-change mRNA levels (Eq 6) for the TF Gt at the percent embryo length indicated. The line representing *ρ* = 1 (Eq 7) is indicated with a gray line. Points below this line are robust, while points above are sensitive. (C) The ratio of the variance in fold-change mRNA to the variance of fold change TF concentration *ρ* (Eq 7) for the indicated TF at each position in the embryo. *ρ* values have been square root transformed for better visual presentation. The dashed line indicates the value *ρ* = 1. Perturbation size *A* in Eq (9) was set to 0.1.

In general, we note that expression at interstripes is more sensitive to fluctuating TF levels than expression at stripe peaks (Figs 1C,S7). Each stripe border is particularly sensitive to the transcriptional repressor that is responsible for forming that expression boundary.

### 4.3 The *eve* locus is *r*-robust with respect to nucleotide changes

Genetic systems may also be robust with respect to changes in DNA sequence. In order to investigate the robustness of the *eve* locus with respect to sequence perturbation, we simulated random mutations to *r* nucleotides 10,000 times, with *r* spanning 1 to 10% of all nucleotides (see Materials and Methods Section 5.3). This results in a curve that saturates with increasing *r* (Fig 2A). This saturating curve is well described by a system with a limited number of sensitive parameters. When we fit Eq 3 at every position along the anterior-posterior axis, we find that stripe peaks are more robust to perturbation than interstripes in that they have fewer sensitive parameters (Fig 2B).

**Figure 2.**
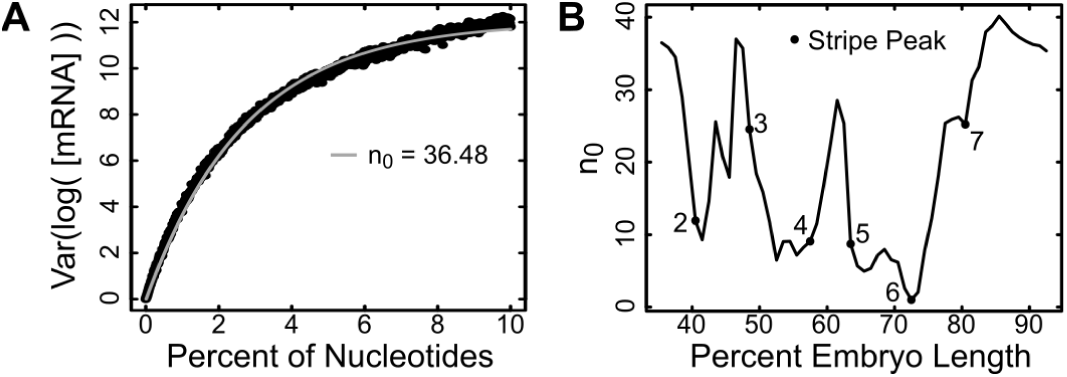
Robustness of *eve* expression to mutation of DNA sequence. (A) The variation of *eve* expression at 35.5% embryo length at various values of *r* nucleotides that are mutated. The fit to data (Materials and Methods) is shown as a grey line, and the estimated number of sensitive nucleotides, *n*_0_, is indicated (Eq 3). (B) The number of sensitive nucleotides, *n*_0_ at every position along the anterior-posterior axis.

### 4.4 Longer *eve* enhancers are more robust to perturbation

For the second stripe of *eve*, four sequences of different length are known to drive expression: the intact locus, the proximal 1700 bp, S2E, and MSE2. Each larger sequence contains the sequence of all smaller enhancers (Fig 3A). In order to investigate whether additional sequence contributes additional robustness, we investigated the number of sensitive nucleotides *n*_0_ in each of these four sequences at the peak of stripe 2 expression (40.5% embryo length). We find that as sequence length grows, not only does the ratio of sensitive nucleotides decrease, the absolute number of sensitive nucleotides decreases from about 26 to 12 (Fig 4A-D).

**Figure 3.**
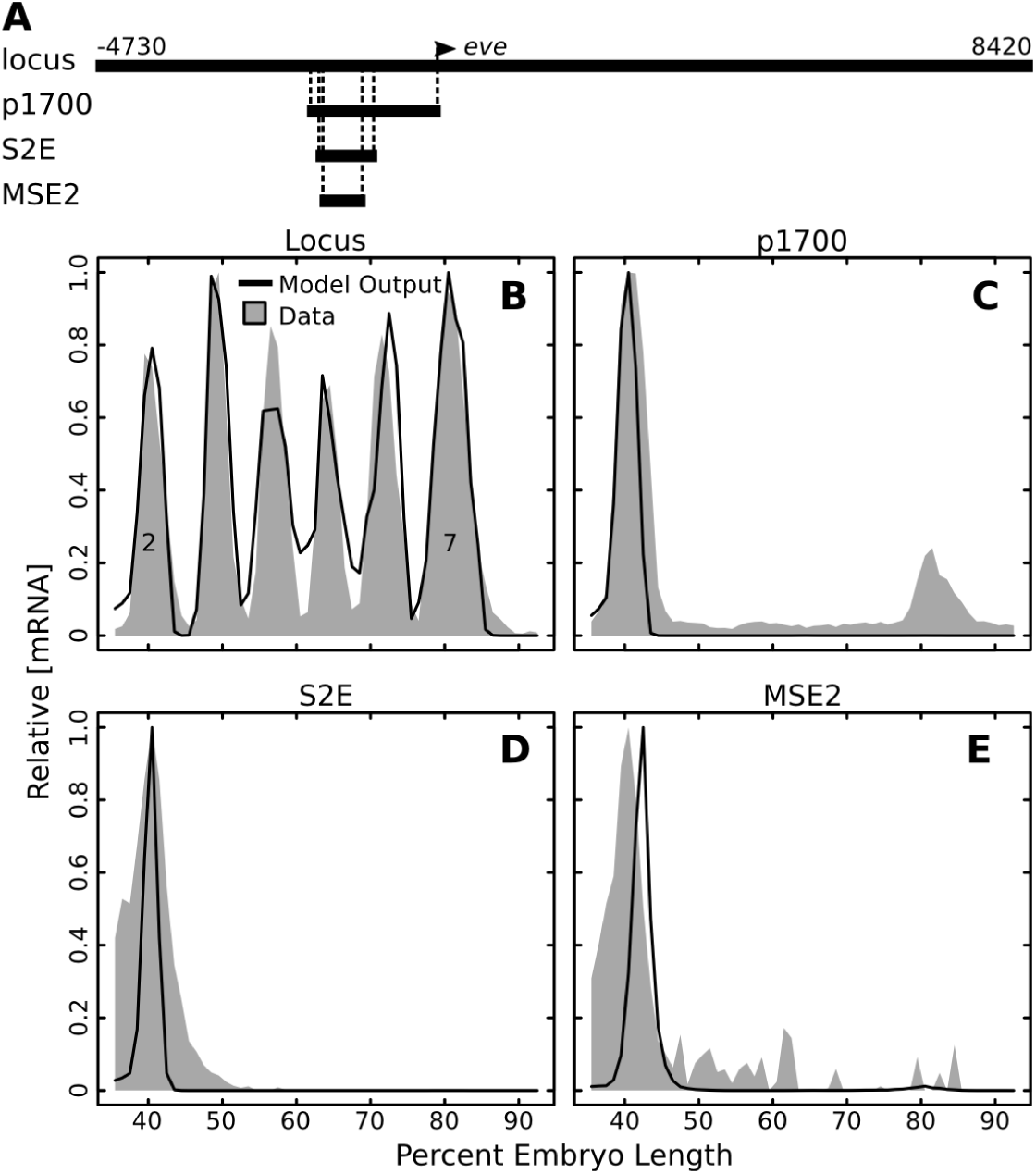
Predicted expression driven by successively smaller enhancers. (A) Diagram showing the entire *eve* locus and successively smaller sequences that all drive stripe 2. Where each sequence aligns within the locus is indicated with dashed lines. Position with respect to TSS is indicated. (B-E) The model predicted expression and actual expression for each of the sequences in (A) along the anterior-posterior axis. Model output is in black lines and expression data is in gray shading. Relative expression on a 0 to 1 scale is reported.

**Figure 4.**
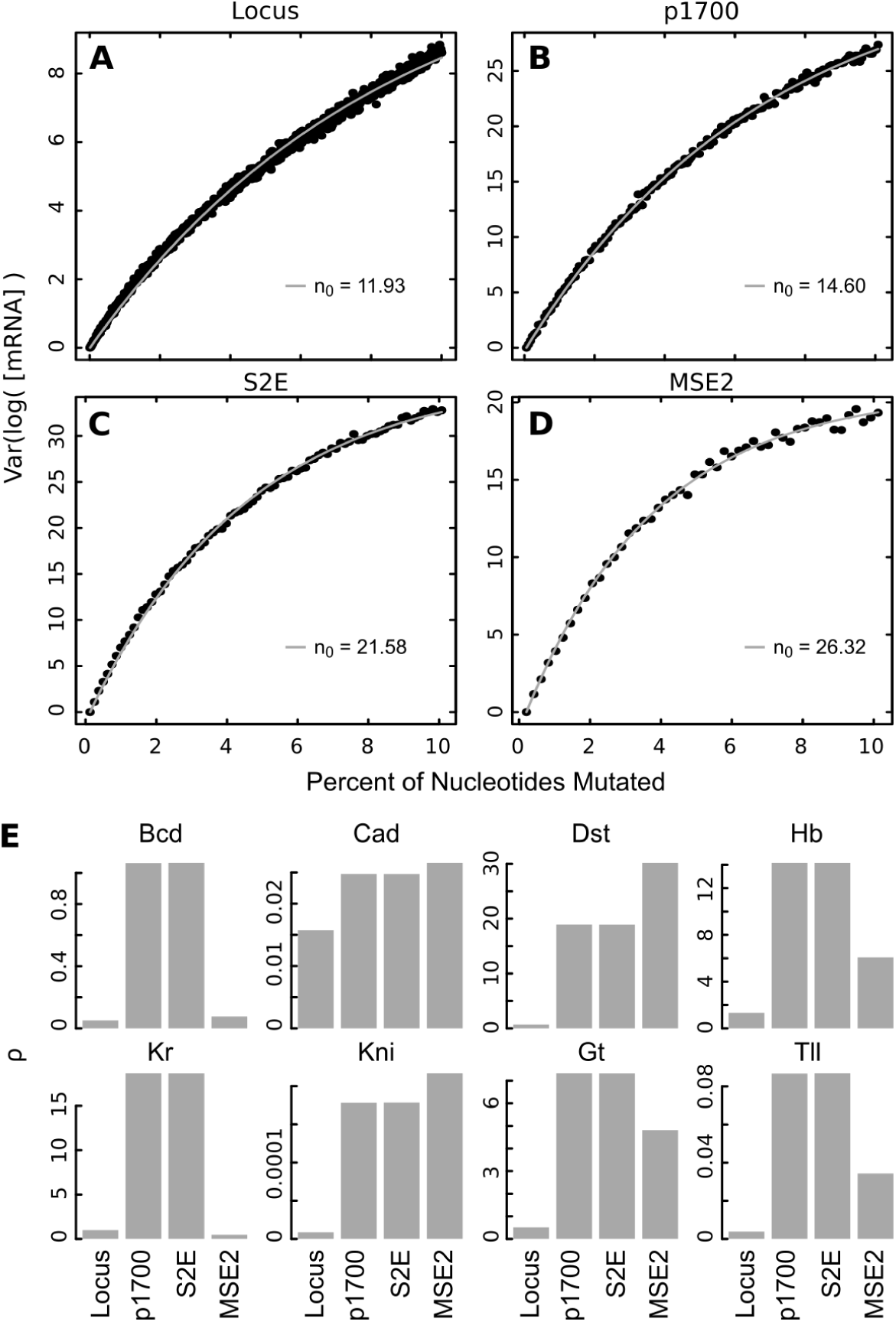
Mutational robustness of natural S2Es. (A-D) The variation in mRNA expression for increasing numbers of perturbed nucleotides at 40.5% E.L. (peak of stripe 2) is shown for sequences of different length that drive *eve* stripe 2. The best fit curve and estimated number of sensitive nucleotides *n*_0_ is indicated. (E) The ratio of variation in mRNA to variation in TF concentration *ρ* (Eq 7) at 40.5% E.L. for each TF and each sequence. *A* was set to *A* = 0.1 in Eq (9) for simulations.

We also investigated robustness to changes in TF concentration for each of the stripe 2 enhancers. For the majority of TFs, there was a relationship between the size of the enhancer and robustness to TF concentration (Fig 4E). In all cases, the intact locus was the most robust to changes in TF concentration. With respect to some TFs, MSE2 was more robust than S2E or the proximal 1700bp.

### 4.5 Robustness is a function of enhancer length

In order to determine whether the relationship between sequence length and robustness to mutation is inherent to the system, we generated 8010 putative stripe 2 elements *in silico*. We generated 10 putative S2Es with each length from 200 bp to 1000 bp. All 8010 S2Es are predicted to drive the correct expression pattern (Fig 5A). We estimated the number of sensitive nucleotides *n*_0_ for each of these S2Es by simulating 10,000 sets of *r* sequence mutation with *r* spanning from 1 bp to 10% of the sequence length. We estimated *n*_0_ at the peak of stripe 2 expression (40.5% embryo length). We found that *n*_0_ decreases with enhancer length (Fig 5B).

**Figure 5.**
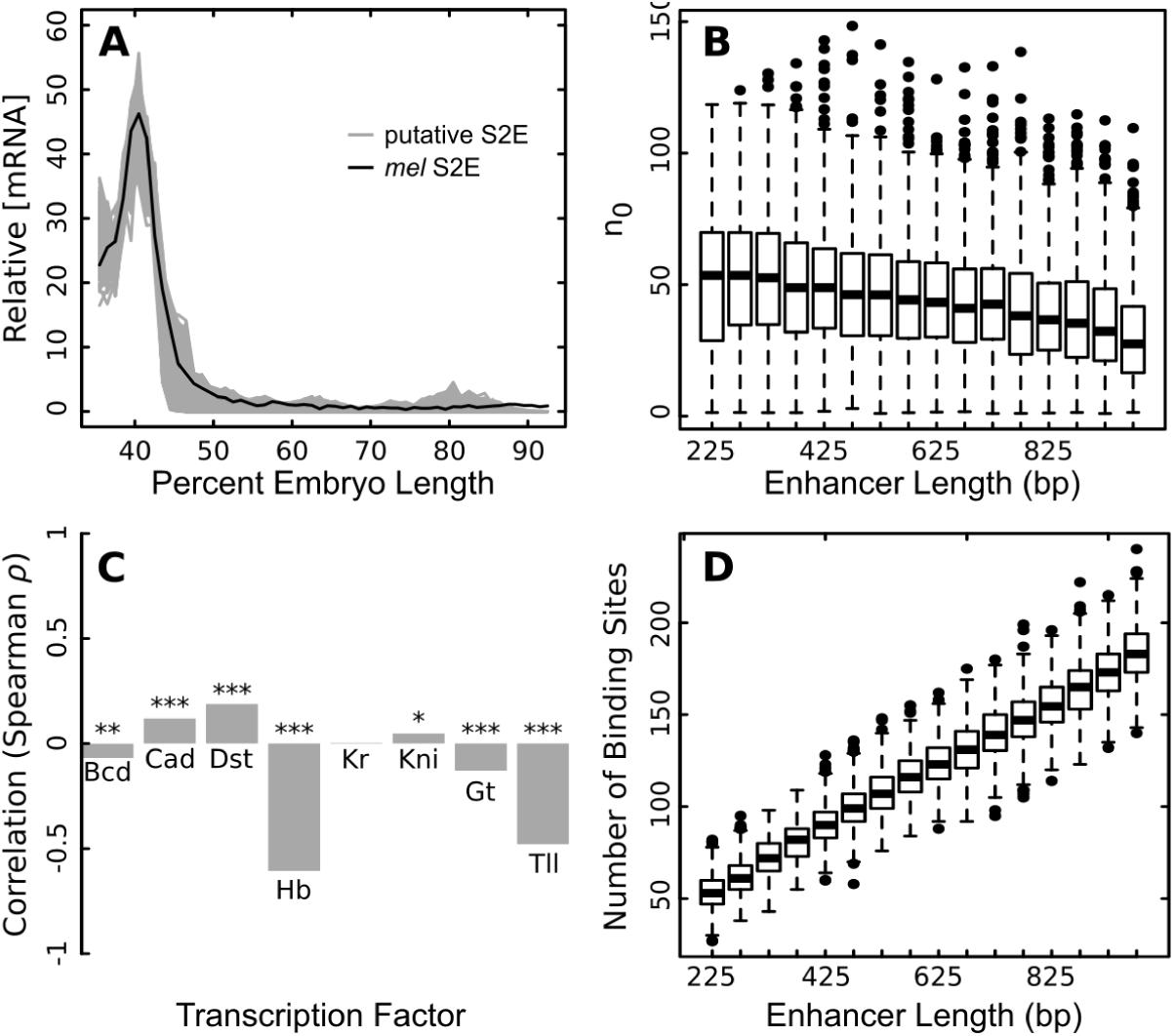
Sequence robustness of putative S2Es. (A) Predicted expression of 8010 putative S2Es with lengths from 200 bp to 1000 bp (10 each, gray lines). Each putative S2E is predicted to drive expression similar to the natural *D. melanogaster* S2E. (B) Boxplots of the number of sensitive nucleotides *n*_0_ *vs.* sequence length for each of the 8010 putative S2Es in bins of 50 bp. *n*_0_ is significantly correlated with sequence length (Spearman *ρ* = −0.25, *p* < 2.2 ×10^*-*16^). (C) The correlation (Spearman *ρ*) between the sensitivity to changes in transcription factor concentration (*ρ*) and the length of putative S2Es (* *p* < 10^*-*4^; ** *p* < 10^*-*9^; *** *p* < 10^*-*16^). The ratio of the variance in mRNA to the variance in TF levels, *ρ* (Eq 7) was estimated with *A* = 0.1. (D) Boxplots of the number of modeled binding sites in each putative S2E *vs.* sequence length for each of the 8010 putative S2Es in 50 bp bins. Spearman *ρ* = 0.96, *p* < 2.2 × 10^*-*16^.

We also investigated robustness to changes in TF concentration for each of these 8010 enhancers. We measured *ρ* from 10,000 simulations with *A* = 0.1 at 40.5% embryo length. For most TFs, there is a relationship between enhancer length and robustness to changes TF concentration, but the direction of this relationship was not consistent between factors (Fig 5C).

### 4.6 The location and mechanism of sensitive nucleotides

In order to identify the location of sensitive nucleotides we tested all possible single nucleotide sequence perturbations and selected the nucleotides that led to the highest log variance of mRNA expression (Fig 6A). For all sequences except MSE2, the sensitive nucleotides occurred in a tight cluster at the 3’ end of the stripe 2 enhancer.

**Figure 6.**
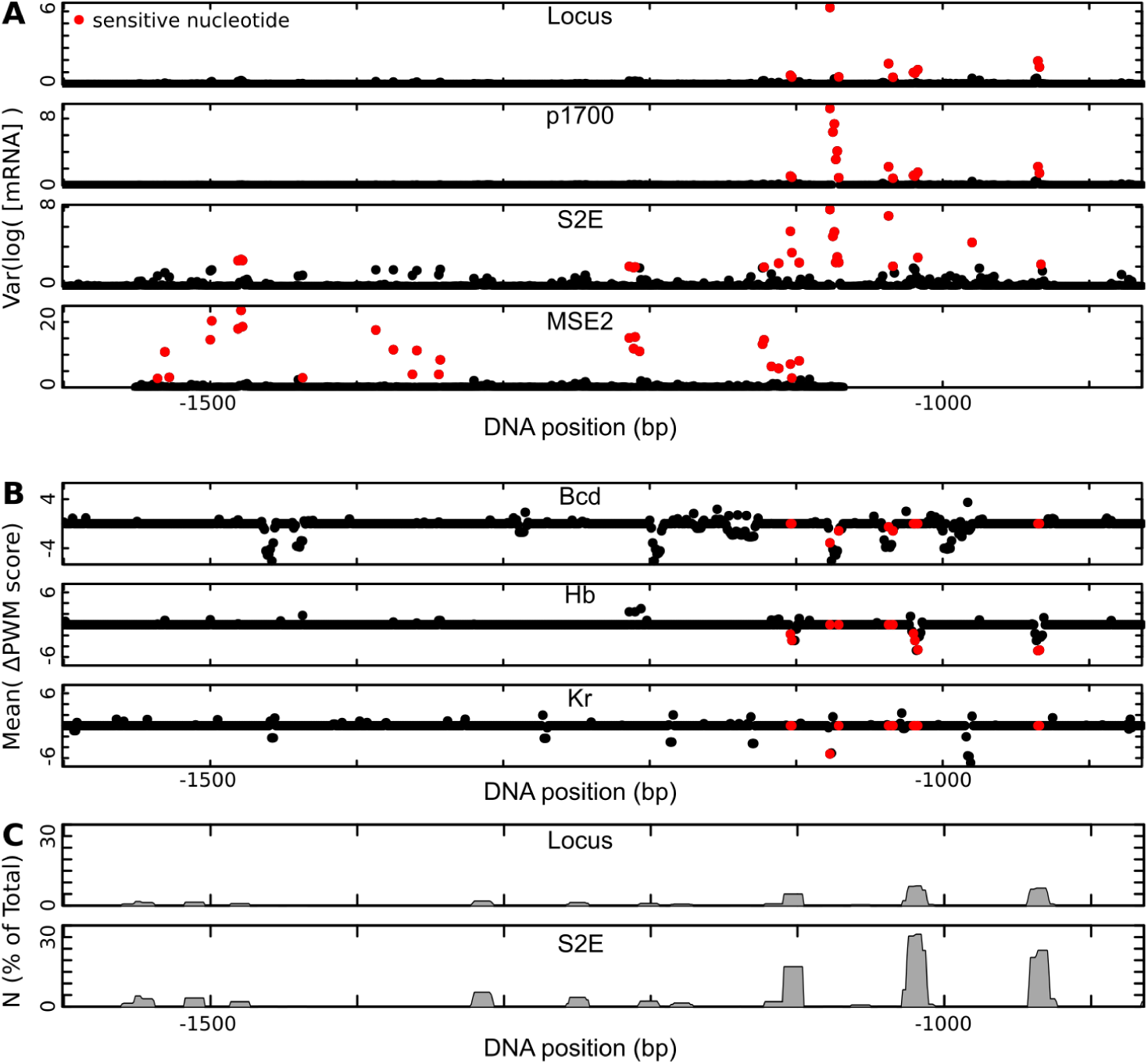
Location of sensitive nuclceotides. In each graph, the horizontal axis is restricted to S2E, and shows sequence position relative to the transcription start site. (A) Log variance of mRNA expression when each nucleotide is perturbed one at a time, for each of the four sequences tested. The *n*_0_ most sensitive nucleotides are indicated in red. (B) The mean change in PWM score for the indicated factors when each nucleotide in S2E is perturbed. The *n*_0_ most sensitive nucleotides are indicated in red. (C) The percent of total strength of transcriptional activation (*N*) along DNA for the locus (top) and S2E (bottom). Sensitive nucleotides represent a smaller percent of total activation in the locus compared to S2E.

**Figure 7.**
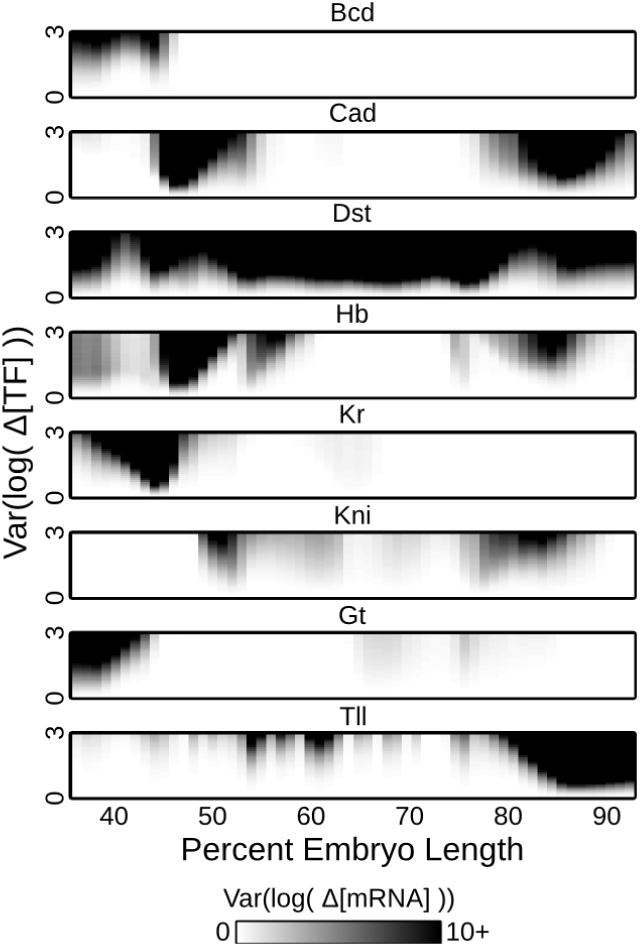
Robustness of *eve* expression to variation in TF concentration. A heatmap comparing the variance in fold-change input to fold-change output (Eqs 6-7) Var(log(Δ[mRNA])) at different positions within the embryo as well as different sizes of perturbation to TF concentration, indicated by Var(log(Δ[TF])). Darker shading represents increasing variation in mRNA levels.

**Figure 8.**
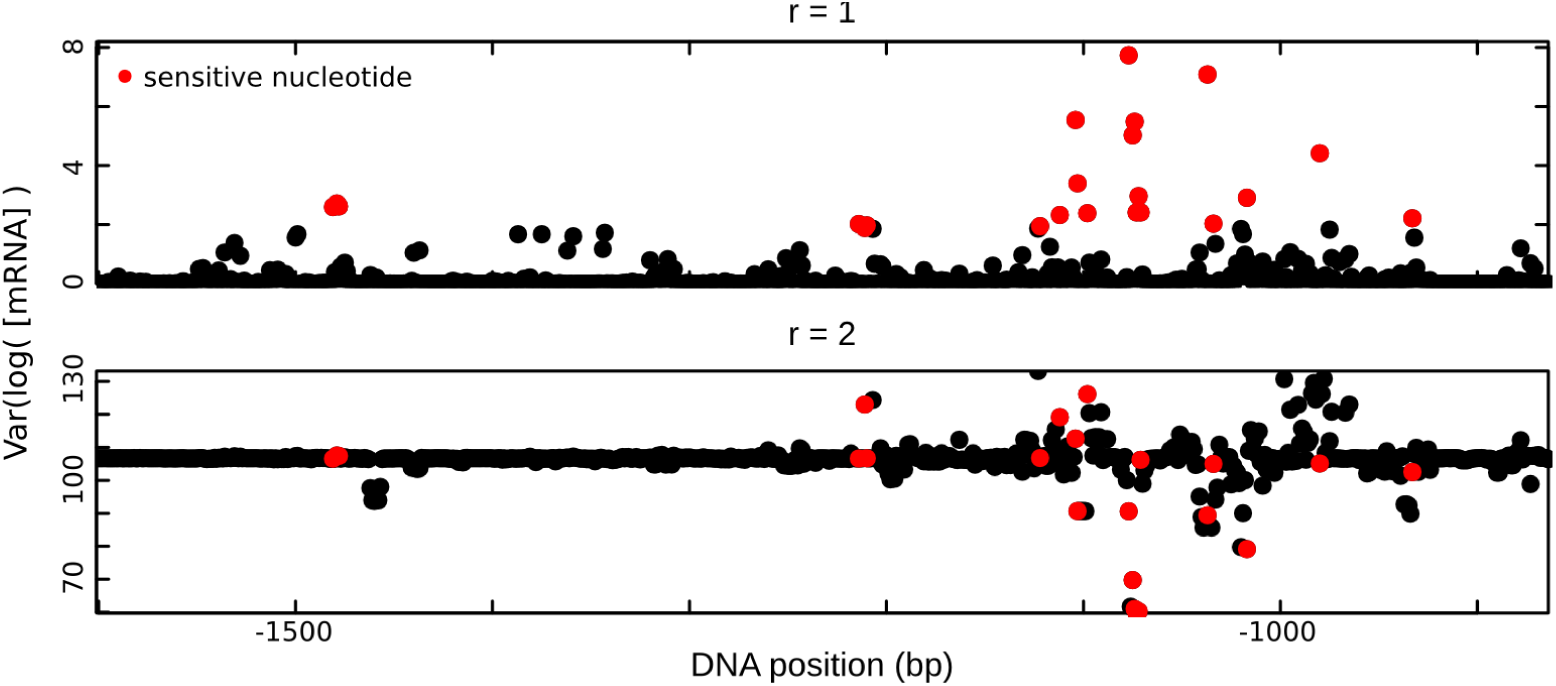
Sensitive nucleotides change with. *r* (Top) The log variance in S2E mRNA expression when each nucleotide is perturbed one at a time (*r* = 1). The top 26 most sensitive nucleotides are indicated in red. (Bottom) The log variance in S2E mRNA expression when each nucleotide is perturbed in a pairwise fashion with all other nucleotides (*r* = 2). The 26 most sensitive nucleotides from the *r* = 1 are indicated in red.

To identify which factors drive sensitivity, we tested the mean change in PWM score for the eight TFs considered here. Sensitive nucleotides tend to lead to a reduced PWM score for at least one of Bcd, Kr, and Hb, but not all binding site losses correspond to sensitive nucleotides (Fig 6B). This effect was especially strong for Hb.

The three Hb sites that contain sensitive nucleotides are responsible for a considerable amount of the total activation in S2E (Fig 6C), however the percent of total activation due to these sequences is much smaller in the intact *eve* locus (Fig 6C). This distributed activation makes the *eve* locus more robust to perturbation of these sequences than smaller sequences.

### 4.7 Sensitive nucleotides are more conserved

Mutations of sensitive nucleotides, which lead to a greater change in transcription rates, are likely to have a negative effect on organismal fitness. Thus, we expect sensitive nucleotides to be more conserved than insensitive nucleotides. To test this hypothesis, we performed an alignment of S2Es from 12 Drosophilids to assess conservation at every nucleotide. Using the sensitivity values from the intact *eve* locus we found that the sensitive nuclei were conserved in 79.5% of species, on average, compared to a background conservation rate of 68%. The correlation between sensitivity and conservation was significant (Spearman *ρ*; *p* = 0.042).

## 5 Discussion

In this work we assessed the robustness of enhancers with respect to changes in TF levels or sequence mutation in the context of a sequence level model of gene regulation. We found that enhancers are *r*-robust to single nucleotide sequence changes, and that this robustness increases with the length of the sequence. The precise level of *r*-robustness seen, however, is not solely dependent on DNA sequence. It depends on the state of bound TFs, and thus manifests itself experimentally as dependence on position within the embryo. Sensitivity, when observed, is coupled to biological function. This point is most clearly seen in the dependence of domain border positions on TF concentration, but is also observable in the functional role of sensitive nucleotides. We discuss each of these points below.

Our major finding is that robustness of *eve* to sequence mutation is well-described by *r*-robustness. That is, the *eve* regulatory DNA is a system with a small number of sensitive parameters. This is understandable in terms of the classic experiments that elucidated this regulation. These experiments showed that enhancer function resided in multiple binding sites for each TF, each of which could be disrupted by site-directed mutation [51, 19]. Such a picture is fully compatible with *r*-robustness with *n*_0_ equal to about 26 for MSE2, which is on the order of the number of the number of base changes required to mutate all of the binding sites for a single TF.

Our finding that *n*_0_ decreases to only 12 for the whole locus indicates that robustness increases with increasing length of regulatory DNA. This might appear to contradict the modular structure of enhancers, but it too is compatible with the experimental literature. Early experiments with enhancers sought to find the minimum fragments that could recapitulate an expression feature using non-quantitative assays [19], MSE2 is quantitatively not equivalent to the full S2E, expressing at a level 5 times lower [41]. Moreover, MSE2 provides a lower rate of rescue of lethality than does the full S2E [53]. S2E, in turn, was first identified by the presence of two conserved sequences at either end [54], but this structural feature says nothing about the actual functional limits of S2E, which are known to be larger in the closely related species *D. erecta* [55]. Moreover there is evidence from the sea squirt *Ciona* that redundancy built into enhancers ensures robust expression in appropriate tissues without disrupting specificity [56]. Redundancy also buffers environmental perturbations, which can disrupt minimal enhancers [53, 57]. We have already alluded to the existence of shadow enhancers, redundant enhancers controlling the same expression domain and thereby increasing robustness [21, 20, 22, 23, 58, 24, 59, 25, 26, 60].

We also compared the level of robustness of putative *in silico* stripe 2 enhancers of different lengths, and also found that longer enhancers were more robust. These results also suggest that selective forces that are not explicitly modeled here drive the evolution of robustness. These *in silico* enhancers were selected for their pattern generating capability, but not explicitly for robustness. Thus, it is interesting that both MSE2 and S2E have fewer sensitive nucleotides (26.3 and 21.6) than *in silico* enhancers of the same length (average of 49.5 and 37.1), indicating that robustness to sequence perturbation may be selected for in natural populations, a conclusion reinforced by our finding that sensitive nucleotides are better conserved across species. The high correlation between length and number of sites (Fig 5D) indicates that redundancy in binding sites combined with a more distributed contribution of all the nucleotides, with sensitive nucleotides being responsible of a lower percentage of the total activation, is probably the cause of increased robustness. Supporting this, we find that MSE2 and S2E have a higher density of sites (114 and 204) than the *in silico* enhancers of the same length (average of 100.6 and 141).

It is important to note that all these measures of sequence robustness varied according to position in the embryo. This is natural and expected because the transcriptional state is dependent not only on position, but also on bound TFs, which vary by position. As a consequence, assays for sequence and gene product concentrations provide a very incomplete picture in the absence of knowledge of state information about regulators.

In contrast with the robustness that we see in *cis-*regulatory sequence, we find marked sensitivity to TF concentrations. For example, Fig 1 shows levels of *ρ* exceeding 100. Specifically, the transcription rate at stripe borders was drastically sensitive to changes in the concentration of TFs. For instance, the anterior border of *eve* stripe 2 is controlled by Gt [19]. When Gt levels fluctuate at this embryo position, transcription levels fluctuate to a greater degree. Such sensitivity may be a necessary feature of the circuit, where high sensitivity to individual repressors allows the formation of extremely sharp borders. Figure 1 also shows areas of low sensitivity. Such areas of apparent robustness fall into two classes. First, in Figure 1C there are areas of high robustness in areas where a transcription factor is not expressed. These areas include the region posterior to 50% E.L. for Bcd, Hb between 60% and 70% E.L., Kr posterior to 70% E.L., Kni anterior to 45% E.L., Gt from 45% to 60% E.L., and Tll anterior to 80%. Here the insensitivity to perturbation arises from the trivial reason that any multiplicative factor applied to zero gives zero. Second, we see reduced sensitivity at stripe peaks. We believe this arises because of the dependence of the transcription activation on the individual transcription factors concentration is a strongly nonlinear sigmoidal function whose derivative is very large close to a threshold (domain borders) and low far away. This excludes robustness at the domain borders and allows it in between.

The results reported here are a precise and quantitative characterization of what it means for a specific biological system to be “robust but fragile.” The concept of *r*-robustness makes this idea precise, and in the case of robustness against changes of sequence, the values of *n*_0_ found have a clear relationship to well known experimental results. We see no evidence of the 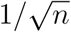 concentration characteristic of the central limit theorem as illustrated in Eqs. 4 and 5. This is to be expected in biological systems, where the necessity of precise control confronts the need for stability and resilience against perturbation, a concept well captured by the idea of *r*-robustness.

The *r*-robustness of *cis-*regulatory sequences is achieved by a principle of “hierarchical heterogeneity” according to which nucleotide mutations have widely distributed impacts. The total rate of transcription of a gene is a weighted sum of contributions of different DNA segments. The contribution to the total transcription of each segment is itself a sum over many interacting sites. At each level of this hierarchy, sensitive functions such as exponential binding affinities or the diffusion-limited Arrhenius rate law produce a disparity among parameters, in which sensitive parameters stand out. Sensitive nucleotides dominate the others and their impact cannot be easily flattened by addition, thus ousting distributed concentration effects. However, in longer sequences the combined effects of many nucleotides dilutes the impact of all the mutations. Therefore, some nucleotides, sensitive in short sequences, become insensitive in the long ones.

## 6 Materials and Methods

### 6.1 Model selection

The model used in this work is the same as reported in Barr and Reinitz[41]. The parameter set used was the best model including chromatin state information, called ‘Repeat Chromatin #2’ in that work.

### 6.2 Simulations of TF concentration perturbation

We perturbed TF concentration by selecting a random number *X* distributed uniformly between −1 and 1. Then we multiplied the TF concentration at every embryonic position by exp(*AX*), where *A* is a set parameter that scales the size of perturbation, and we observed the predicted change in mRNA synthesis rate at all positions. We repeated this calculation 10,000 times for each value of *A* between 0 and 3 in increments of 0.1, for a total of 310,000 simulations. This was repeated for each of the 8 TFs included in the model.

### 6.3 Simulations of sequence mutation

In the intact locus model, some nucleotides are not accessible for TF binding because of the chromatin state. We perturbed DNA sequence by selecting sets of *r* nucleotides only from open chromatin regions. The *r* nucleotides were then substituted by one of the remaining three possible nucleotides with equal probability. We then assessed model output at each position. This was repeated for every set *r* from 1 10% of the total accessible nucleotides.

### 6.4 Estimation of sensitive nucleotides

To fit Eq (3) to data, we used simulated annealing from the R package GenSA, using default parameters. *n* was set to the total length of accessible nucleotides, or 8765 for the locus, 1726 for p1700, 804 for S2E and 484 for MSE2.

### 6.5 Enhancer-reporter assays

The locus expression, as well as the expression of the S2E and MSE2 construct are reported in Barr and Reinitz[41]. The p1700 data is from Janssens *et al.*[33].

### 6.6 Generation of putative S2Es

To generate putative S2Es of different lengths, we fixed the kinetic parameters and optimized DNA sequence using previously described methods[61]. We used the expression of S2E as the target. We started each optimization with a random sequence of the desired final length.

### 6.7 12 species alignment

To identify conservation at sensitive nucleotides we first obtained putative S2E sequences by using the BLAST tool at FlyBase[62]. We identified significant contiguous alignments for the species *sim, sec, ere, yak, rho, ele, tak, eug, bia, kik*, and *pse*. We performed an alignment using Clustal Omega[63] using default parameters. To get a conservation score at every base in the *melanogaster* sequence, we calculated the percent of species containing the same nucleotide as *melanogaster* at that position.

## 7 Acknowledgments

This work was supported by an award from the “France and Chicago Collaborating in the Sciences” (FACCTS) and by grant R01 OD010936 from the U.S. National Institutes of Health.

## 7.1 Appendix

The sequence level model of gene regulation used in this work was the subject of prior investigation[41]. This calculation has two major sections. In the first, chemical rules are used to define the equilibrium binding of transcription factors (TFs) on DNA. In the second, phenomenological rules—which restate important experimental results in the form of carefully selected equations—are used to compute the effects of DNA-bound transcription factors on local gene expression. The full calculation includes six steps: (1) calculation of TF binding affinity to DNA using position weight matrices (PWMs), (2) calculation of fractional occupancy at each binding site identified in step 1, (3) coactivation of Hunchback by locally bound Bicoid or Caudal, (4) quenching of activators by locally bound repressors, (5) weighted summation of unquenched activators along DNA, and (6) transcriptional activation though a diffusion limited Arrhenius rate law. In the sections that follow, we introduce the equations for each of these steps.

### 7.1.1 PWM scores

First, we calculate PWM scores along DNA. We index binding sites with index *i* and TFs with index *a*. We represent the binding site *i* for TF *a*, spanning nucleotides *m* through *n* using the notation *i*[*m, n*; *a*]. When it is necessary to discuss the positioning and possible overlap of binding sites, we denote by *m*_*i*_, *n*_*i*_, *a*_*i*_ the leftmost and the rightmost positions of nucleotides (measured on the DNA in the 5′to 3′direction) spanning the binding site *i*, and the TF binding the site *i*, respectively. The binding site indices *i* are ordered such that *m*_*i*_ < *m*_*j*_ for any *i* < *j*. If *n*_*i*_ ≥ *m*_*j*_ the sites *i, j, i* < *j* overlap and are competing.

The PWM score *S* for each binding site *i* of the TF *a* is given by

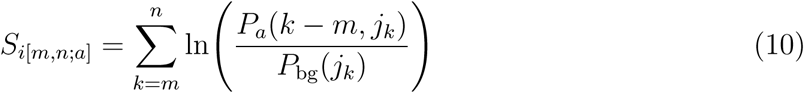

where *j*_*k*_ ∈ *{A, C, T, G}* is the nucleotide observed at position *k, P*_*a*_(*k-m, j*) is the probability of observing this nucleotide at position *k - m* in a binding site for factor *a*, and *P*_bg_(*j*_*k*_) is the probability of observing this nucleotide in the null distribution, which we take to be the *D. melanogaster* nucleotide frequencies: *P*_bg_(*C*) = *P*_bg_(*G*) =0.203; *P*_bg_(*A*) = *P*_bg_(*T*) = 0.297. We only keep binding sites with positive scores, representing sites that are more likely to be binding sites than the background distribution.

The PWM score *S* is used to calculate the binding affinity *K* [64].

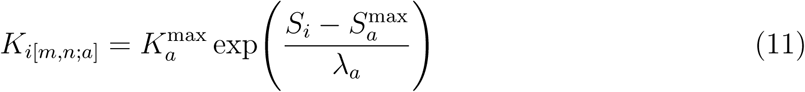

where 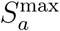 is the maximum score possible for the PWM of factor *a* and *λ*_*a*_ is a positive proportionality constant.

### 7.1.2 Adjustment for non-specific binding affinity

Binding affinity has both sequence specific and non-specific contributions. We only want to consider specifically-bound TFs in our calculations. Previous work has shown that non-specific binding energy is approximately three orders of magnitude smaller than the maximum specific binding energy[65]. Following the derivation described [41], we calculate an effective binding affinity,

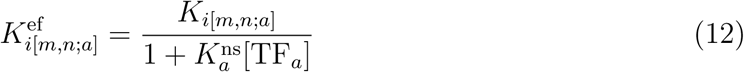

where *K*^ns^ is the non-specific binding energy of TF *a*, which we fix to 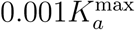.

### 7.1.3 Fractional Occupancy

The fractional occupancy represents the portion of time any particular binding site *i* is bound by transcription factor *a*. This can be computed using from the transcription factor concentration and the binding affinity given in Eq 12. To calculate fractional occupancy, we add the Boltzmann weight of all binding states that contain a bound transcription factor, divided by the sum of the Boltzmann weights of all the binding states, known as the partition function. We take into account the cooperativity of binding of close sites and the competition for binding between overlapping sites.

To this aim we introduce *q*_*i*_, 0 ≤ *q*_*i*_, representing the Boltzmann weight of the bound state of the site *i*. The weight *q*_*i*_ is proportional to the product between the effective binding affinity and the concentration of the transcription factor, giving

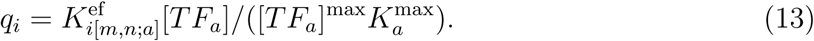

We also define a function that specifies the index of the rightmost (closest to 5^*t*^) non-competing binding site to the left (in the direction towards 5^′^) of the site *i*:

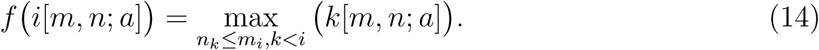

If there is no non-competing binding site left of the site *i*, then *f* (*i*[*m, n*; *a*]) = 0.

We define iteratively the following three partial partition functions:

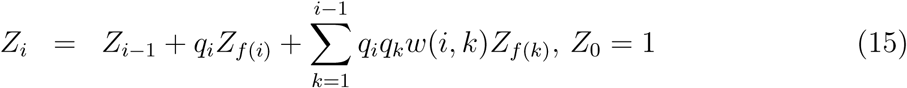

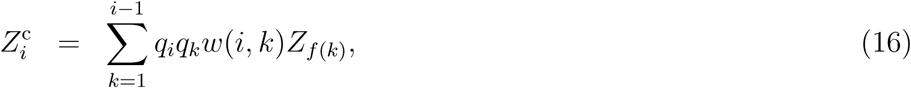

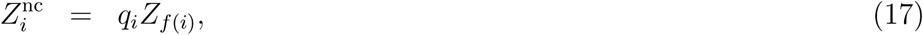

where *w*(*i, k*), with 0 ≤ *w*(*i, k*) ≤ 1, is the cooperative interaction strength between sites *i* and *k. w*(*i, k*) is only nonzero for Bcd sites less than 60 bp apart.

We compute the partial partition functions while scanning DNA in the direct (5’ to 3’) and in the reverse (3’ to 5’) directions. We denote these 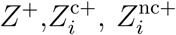, and 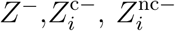, respectively. From these quantities, we compute the fractional occupancy as

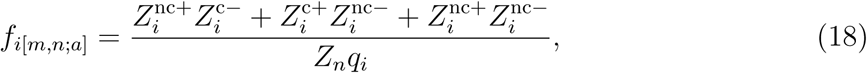

where *n* = max *i*. A more detailed description is given elsewhere [66].

### 7.1.4 Coactivation

For sites *i* and *k*, the distance between sites is computed as

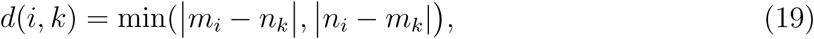

and the efficiency of interaction is given by

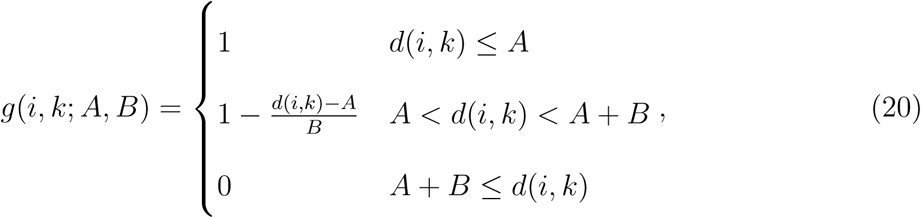

where *A* and *B* are positive parameters governing the shape of this interaction.

We divide fractional occupancy into two states: an activating state *f* ^*A*^ and a quenching state *f* ^*Q*^. For obligate repressors 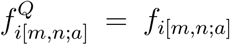, and similarly for obligate activators 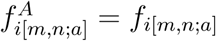. For factors that coactivate,

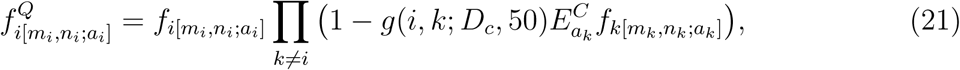

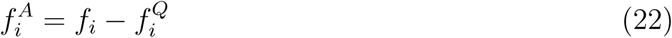

where *D*_*c*_ is a free parameter giving the maximum distance at which coactivation is 100% efficient and 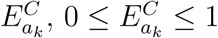, is a free parameter giving the maximum efficiency with which the factor *a*_*k*_ induces activation of factor *a*_*i*_. This product occurs over all *k* binding sites within the locus.

### 7.1.5 Quenching

The effective occupancy of each activator, F, is given by

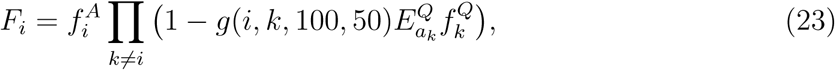

Where 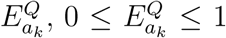, is a free parameter giving the efficiency with which the factor *a*_*k*_ quenches. This product occurs over all *k* binding sites within the locus.

### 7.1.6 Summation of activating strength

The contribution to the total transcriptional activation for a sequence of bases from *m* to *m* + *α* is

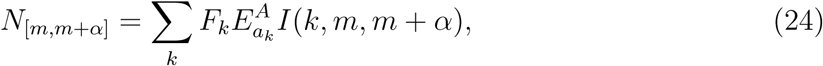

where the 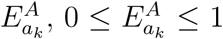 is the activating efficiency of factor *a*_*k*_ and *I*(*k, m, m* + *α*) is a function that specifies whether the site *k* falls between *p* and *q*, given by

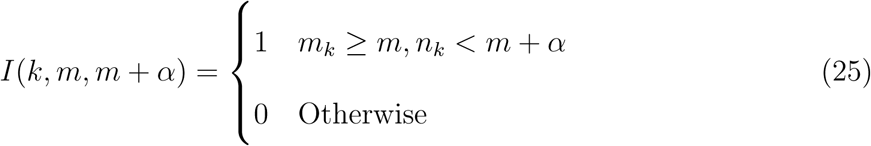

### 7.1.7 Activation by a diffusion-limited Arrhenius rate law

The rate of transcription driven by a sub-sequence bounded by *m* and *m* + *α*, is given by

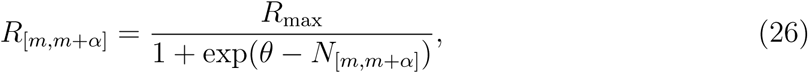

where *R*_max_ ≥ 0 is the efficiency of transcription, *θ* ≥ 1 is the total energy barrier which sets the rate of transcription in the absence of activation. For a locus of length *l*, the fraction of time that any DNA segment [*m, m* + *α*] influences the promoter is given by

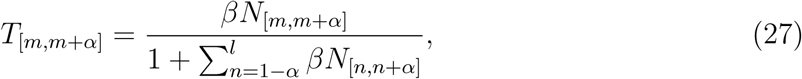

where the free parameter *β, β* ≥ 0, determines how much individual bound adaptors increase the frequency of interaction with the promoter. The total rate of transcription driven by the locus is then given by the frequency-weighted sum of transcription due to each DNA segment [*m, m* + *α*], so that

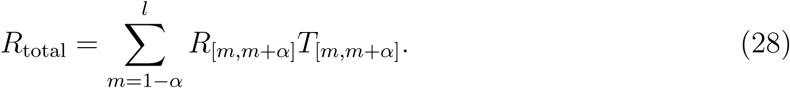

